# Genetically encoded multimeric tags for intracellular protein localisation in cryo-EM

**DOI:** 10.1101/2022.12.10.519870

**Authors:** Herman KH Fung, Yuki Hayashi, Veijo T Salo, Anastasiia Babenko, Ievgeniia Zagoriy, Andreas Brunner, Jan Ellenberg, Christoph W Müller, Sara Cuylen-Haering, Julia Mahamid

**Affiliations:** Structural and Computational Biology Unit, European Molecular Biology Laboratory (EMBL), Heidelberg, Germany; Cell Biology and Biophysics Unit, EMBL, Heidelberg, Germany; University of Heidelberg, Heidelberg, Germany; Collaboration for Joint PhD Degree between EMBL and Heidelberg University, Faculty of Biosciences, Heidelberg, Germany

## Abstract

Cryo-electron tomography is a powerful label-free tool for visualizing biomolecules in their native cellular context at molecular resolution. However, the precise localisation of biomolecules of interest in the tomographic volumes is challenging. Here, we present a tagging strategy for intracellular protein localisation based on genetically encoded multimeric particles (GEMs). We show the applicability of drug-controlled GEM labelling of endogenous proteins in cryo-electron tomography and cryo-correlative fluorescence imaging in human cells.

## Main text

The localisation and identification of macromolecules in crowded cellular landscapes visualised by cryoelectron tomography (cryo-ET) are challenging tasks. Current solutions to the problem include: (1) cryo-correlative light and electron microscopy (CLEM), whereby fluorescence is used to guide cryo-ET sample preparation, image acquisition and interpretation, albeit with localisation errors due to limited resolution in fluorescence imaging and sample deformation during preparation or transfer^1–4^; (2) pattern recognition, by template matching^5^ or convolutional neural networks (CNNs)^6,7^, is generally applicable to large macromolecular complexes with an identifiable structure. Molecular tags with a unique size, shape or density have been suggested as a complementary solution for direct macromolecular localisation in cryoelectron tomograms. However, existing strategies, such as fusion with iron-enriching ferritin^8^ and surface labelling with DNA origami^9^ have limited applicability in mammalian cells due to potential cytotoxicity and limited accessibility beyond the cell surface, respectively. Here, we report on the development of a general tag for intracellular protein localisation in cryo-ET imaging of mammalian cells.

Our design concept encompasses a genetically-encoded tag that is structurally distinct in the cell, easy to recognise in tomograms, and that tethers to GFP (green fluorescent protein) upon addition of a small molecule, thereby enabling labelling of GFP-fusion proteins with minimal interference to protein function (Fig. 1a). To provide a distinct structural signature, we based this tag on an encapsulin protein scaffold. Encapsulins are proteins in archaea and bacteria that form icosahedral particles of defined stoichiometries, ranging from 25 to 42 nm in size^10^. The larger encapsulins have been shown to self-assemble inside mammalian cells and have a distinct appearance in tomograms^11^. Thus, we reasoned that an encapsulin-derived genetically encoded multimeric particle (GEM) would provide a suitable platform for the development of an intracellular tag. By decorating the surface of the encapsulin scaffold with the FKBP-rapamycin-binding (FRB) domain of mTOR, we enable rapamycin-inducible coupling to GFP through an adaptor protein consisting of FKBP and an anti-GFP nanobody^12^. To enable tuning of expression and systematic optimisation of tagging efficiency, GEMs and adaptor proteins are expressed via a single doxycycline-inducible gene cassette and can be fluorescently labelled via a Halo- and SNAP-tag, respectively.

**Fig. 1:**
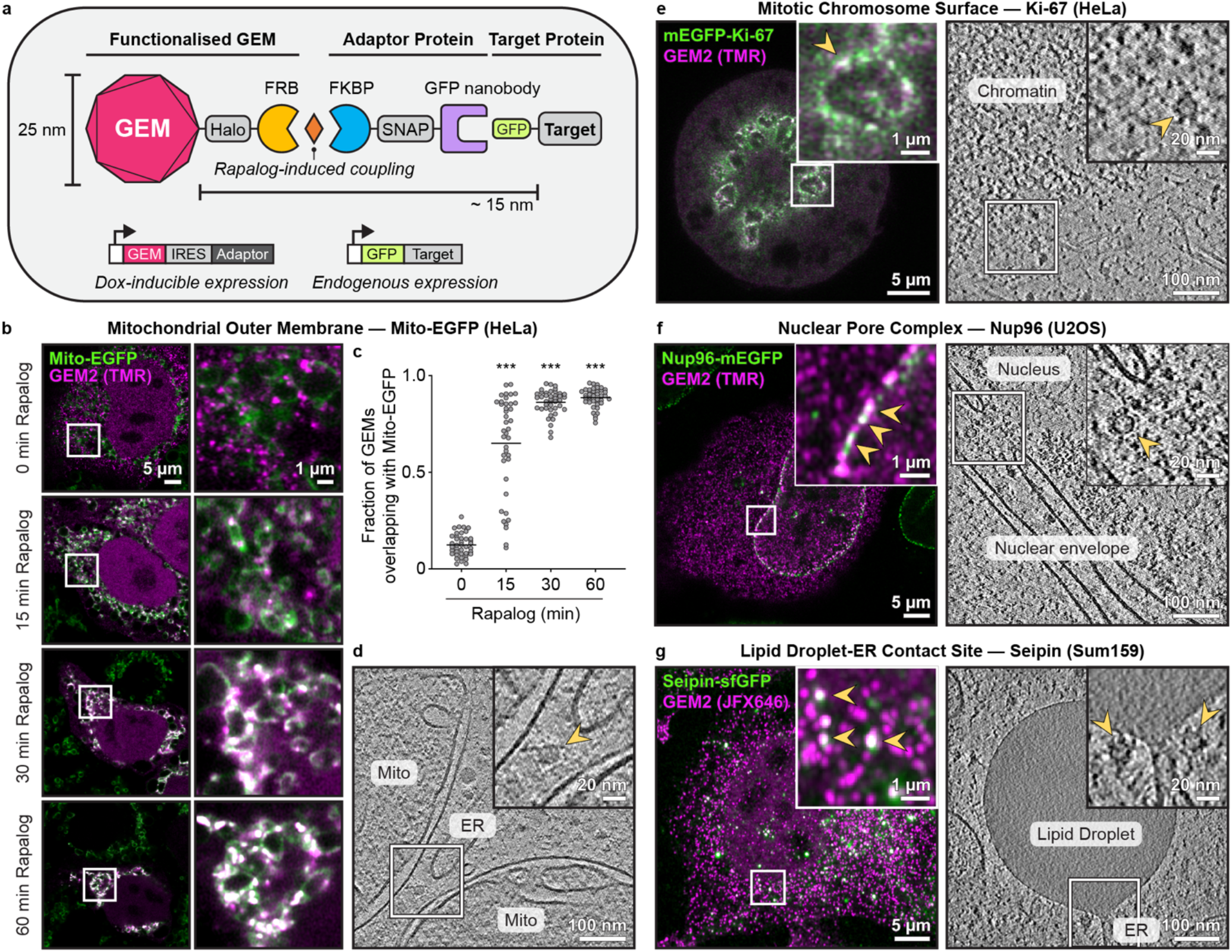
GEM labelling of different GFP-fusion proteins in human cells. **a**, Schematic of the system. **b– c**, Time course of GEM2 (fluorescently labelled with TMR, tetramethylrhodamine; magenta) recruitment to Mito-EGFP (green) upon rapalog treatment by fluorescence microscopy in HeLa cells. **c**, Fraction of GEMs overlapping with Mito-EGFP per cell. Lines indicate mean, n = 40–41 cells per treatment, 2 experiments. P***<0.0005, Kruskall-Wallis test followed by Dunn’s test, compared to 0 min treatment. **d**, Tomographic slice showing GEM2-labelled mitochondria, after 30 min rapalog treatment. Arrowhead in inset indicates a GEM2 particle. **e–g**, GEM labelling of endogenously GFP-tagged proteins in different human cell lines. Left, fluorescence images. Arrowheads indicate examples of colocalization between GEM2 and the target protein. Right, tomographic slices. Arrowheads indicate GEM particles. For the images shown, cells were treated with rapalog for the following durations: Ki-67, 1 h; Nup96, 12 h (fluorescence image) and 8 h (tomogram); seipin, 5 h (fluorescence image) and 10 h (tomogram). The GEM2 and adaptor protein gene cassette was introduced into the AAVS1 locus for expression in the case of Mito, Ki-67 and seipin. For Nup96, the gene cassette was transiently transfected. GEM and adaptor protein expression was induced by 24–48 h of doxycycline treatment prior to rapalog treatment.

To identify a GEM construct compatible with this design, we conducted a library screen in HeLa cells of 10 encapsulins and 3 synthetically designed protein cages, all expected to form 25-nm-sized particles (Extended Data Table 1, Extended Data Fig. 1). Fluorescence imaging of Halo-FRB-tagged constructs revealed 5 candidates that form uniformly sized puncta in the cytoplasm (Extended Data Fig. 2). Among them, the functionalized *Synechococcus elongatus* Srp1 encapsulin (GEM2) also localises to the nucleus. We then evaluated the candidates’ potential for coupling to GFP upon rapamycin treatment using cells stably expressing EGFP on the mitochondrial surface as a test case. We found that GEM2 colocalized with mitochondrial-targeted EGFP (Mito-EGFP) most efficiently (Extended Data Fig. 3). We further confirmed that GEM2 was mobile in the cytoplasm (Extended Data Fig. 4, Supplementary Video 1), and could be recruited to Mito-EGFP within 15 min upon rapalog or rapamycin treatment (Fig. 1b,c, Supplementary Video 2). In agreement with these observations, cryo-ET imaging of focused ion beam (FIB) lamellae from cells after 30 min of rapalog treatment revealed clear icosahedral particles close to the mitochondrial surface (Fig. 1d).

To assess the applicability of the GEM labelling strategy to endogenous human proteins, we targeted GEM2 to three endogenously GFP-tagged proteins at different subcellular locations: Ki-67 at the mitotic chromosome surface^13^, Nup96 at the nuclear pore^14^, and seipin at endoplasmic reticulum-lipid droplet (ER-LD) contact sites^15^. In all cases, we observed rapalog-inducible colocalization of GEMs with the target by fluorescence microscopy (Fig. 1e–g). We note that GEM recruitment dynamics correlated with target protein abundance, as measured by fluorescence correlation spectroscopy (FCS) calibrated imaging^16^: For the more abundant Ki-67, recruitment was rapid and saturated within 30–60 min of rapalog induction, whereas for the less abundant seipin, recruitment continued to increase after 5 h (Extended Data Fig. 5). Further, while a high GEM abundance in the cell gave rise to more complete labelling of the target, it also resulted in more GEMs not bound to the target (Extended Data Fig. 5c, e). This highlights the need to optimise labelling durations and GEM expression levels for different intracellular targets. While the size and multimeric nature of GEMs may be a concern due to steric hindrance and their potential to induce protein target clustering or aggregation, our functional assays indicate that protein function was preserved in all three test cases (Extended Data Fig. 6).

Cryo-ET imaging of rapalog-treated cells revealed visually detectable icosahedral particles near the expected cellular structure for each target protein (Fig. 1e–g). Leveraging their characteristic shape, we trained a CNN for automated localisation of GEM particles in tomograms (Fig. 2a, Supplementary Video 3). By subtomogram averaging of the detected particles, we confirmed that GEM2 forms an icosahedron of the expected size in cells, superposing well with a previously determined *in vitro* structure of the encapsulin scaffold^17^ (Fig. 2b). Additional densities were observed at the 5-fold vertices of the icosahedron, corresponding to the locations of the engineered Halo- and FRB-tags.

**Fig. 2:**
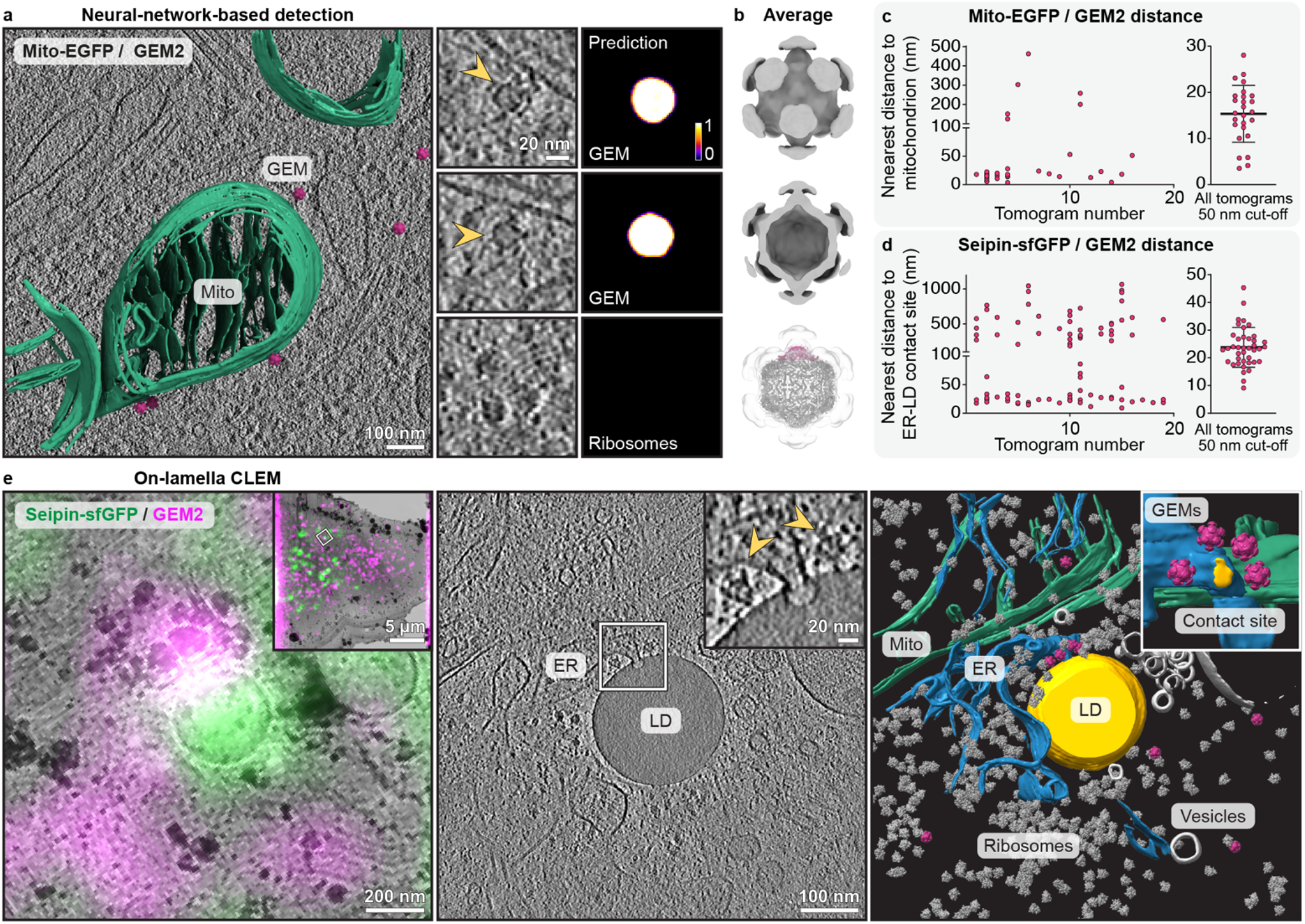
CNN-based detection and multimodal imaging of GEM-labelled proteins. **a**, Tomographic slice of GEM2-labelled Mito-EGFP on mitochondria (see also Supplementary Video 3). Insets show individual GEMs or ribosomes alongside CNN probability scores for GEMs. **b**, In-cell subtomogram average of GEM2 superposed with the structure of the encapsulin scaffold (bottom, PDB 6X8M), with one pentamer coloured in magenta. **c**–**d**, Spatial distributions of GEMs relative to the target per tomogram, n = 34–88 GEMs from 16–19 tomograms. Lines indicate mean ± s.d. in the right panels. **e**, Left, cryo-TEM image of FIB-lamellae overlaid with cryo-Airyscan fluorescence. Middle and right, tomographic slice and corresponding segmentation with insets showing the ER-LD contact site and GEMs. Viewing angle in segmentation inset is rotated to visualize the ER-LD contact site from the LD direction.

Comparing the distribution of GEMs in tomograms of high-abundance and low-abundance intracellular targets, we found that the extent of GEM recruitment correlated with target protein abundance, in line with our fluorescence-based analyses. For the more abundant Mito-EGFP, 76% of particles localised within 50 nm of the mitochondrial surface, whereas for the less abundant seipin, 45% localised within 50 nm of the ER-LD contact site (Fig. 2c–d). We observed a distance of 15 ± 6 nm between the encapsulin surface and the mitochondrial membrane, and a longer distance of 24 ± 7 nm to the ER-LD contact site, likely attributable to the long cytosolic tail of seipin^18^. These measurements provide an estimate of the specificity and precision with which we can localise a target via GEMs.

With 60 Halo-tags providing multiple dye-labelling sites per particle, GEMs offer an additional opportunity for CLEM and an advantage over single GFP-fusion tags. Indeed, upon cryo-fluorescence imaging, we found that GEM fluorescence was sufficiently strong in ∼200-nm-thin FIB lamellae to guide cryo-ET data collection (Fig. 2e). By targeting sites where GEM2 colocalised with seipin, we observed a cluster of GEMs surrounding the target ER-LD contact site, unambiguously revealing the location of seipin and providing evidence for previous hypotheses^15,19^ that the homo-oligomeric seipin encircles the contact site (Fig. 2e). This provides new structural insights into the *in situ* conformation of seipin for a potential mechanism of LD biogenesis.

In summary, we developed a drug-inducible GEM labelling strategy to enable nanoscale-precision localisation of intracellular proteins in cryo-ET. We demonstrate the broad applicability of this tag to GFP-fusion proteins of differing abundances in four subcellular locations in human cells. The modular nature of our design enables further customisation. For example, direct fusion of FRB to the target protein removes the need for an adaptor protein and allows for nanobody-free labelling, with the shorter linker potentially affording more precise localisations (Extended Data Fig. 7). With further development of a more rigid linker or incorporation of asymmetric elements, GEMs may be leveraged to aid subtomogram averaging of structurally-defined macromolecular complexes. Altogether, GEM labelling provides an easy-to-implement strategy for identifying biomolecules in cryo-electron tomograms and thus has the potential to expand applications of cryo-ET to cellular structures where prior structural knowledge is lacking.

## Methods

### Mammalian cell culture

HeLa cell lines were derived from a previously described HeLa Kyoto cell line^20^. HeLa cells endogenously tagged with mEGFP at the N-terminus of Ki-67 were previously described^13^. For generating HeLa cells stably expressing EGFP on the outer mitochondrial membrane, cells were transfected with a plasmid harboring an import signal of yeast mitochondrial outer membrane Tom70p fused to EGFP^21^ using pEGFP-N1 as a backbone (Mito-EGFP) using PEI Max (Polysciences) and cultured for 2 days. Cells were selected using 1 mg/ml G418 (Thermo Fisher Scientific) for 2 weeks and then GFP-positive cells were isolated by fluorescence-activated cell sorting (FACS) using BD FACSAria Fusion, expanded and validated using fluorescence microscopy. HeLa cells were cultured in Dulbecco’s modified medium (DMEM; Sigma) containing 10% (v/v) fetal bovine serum (FBS; Thermo Fisher Scientific), 1% (v/v) penicillin–streptomycin (Sigma), and 1 mM sodium pyruvate (Thermo Fisher Scientific). U2OS cells with knock-in of mEGFP into endogenous locus of Nup96^22^ were cultured in McCoy’s 5A modified medium (McCoy; Thermo Fisher Scientific), supplemented with 10% (v/v) FBS, 1% (v/v) penicillin–streptomycin, 1 mM sodium pyruvate, and 1% MEM nonessential amino acids (Thermo Fisher Scientific). Sum159 cells endogenously tagged with sfGFP at the N-terminus of seipin and seipin knockout cells have been previously described^23,24^. Sum159 cells were maintained in DMEM/F-12 GlutaMAX (Thermo Fisher Scientific) supplemented with 5 mg/ml insulin (Cell Applications), 1 mg/ml hydrocortisone (Sigma), 5% FBS (v/v), 50 mg/ml streptomycin, and 50 U/ml penicillin. All cells were cultured at 37 °C in a 5% CO2-containing atmosphere. These conditions were used for all live cell imaging experiments. All cell lines and plasmids described in this study are listed in Extended Data Tables 2 and 3.

### GEM library screening

Codon-optimised synthetic GEM genes (Biomatik) were subcloned into pcDNA3.1(+) for transient expression under a CMV promoter as Halo-FRB (T2098L) fusions in HeLa cells using PEI Max. Two days after transfection, cells were labelled with 100 nM Halo-TMR (Promega) for 20 min and then 0.2 μg/mL Hoechst 33342 (Thermo Fisher Scientific) for 30 min prior to imaging. For screening of drug-inducible GEM recruitment to GFP, HeLa cells stably expressing Mito-EGFP were transfected with a doxycycline-inducible gene cassette, hereafter the GEM2-Halo-FRB-IRES-adaptor cassette, encoding the GEM-Halo-FRB fusion, an EMCV-derived IRES sequence^25^, and the adaptor protein (FKBP-SNAP-vhhGFP4^12^) under a TREtight promoter^26^. Cells were labelled with 100 nM Halo-TMR ligand and 100 nM SNAP-Cell 647-SiR ligand (New England Biolabs) for 20 min. To induce GEM recruitment to GFP, cells were treated with 0.5 μM rapamycin for 1 h. Screening experiments were performed in live cells on Olympus IXplore SpinSR spinning disk confocal microscope (Olympus) using a 100×, 1.35 NA silicone immersion objective.

### Plasmid construction for CRISPR/Cas9 genome editing

To generate a GEM2 donor plasmid for knock-in into the human adeno-associated virus integration site 1 (AAVS1), Addgene plasmid #129715^27^ was used for Gibson assembly with two gene cassettes: the GEM2-Halo-FRB-IRES-adaptor cassette described above; and rtTA3 transactivator and selection markers separated by 2A self-cleaving peptides (rtTA3-P2A-PuroR-T2A-Thy1.1) under the EF1α promoter. To generate a donor plasmid for knock-in of GEM2-Halo-FKBP in the nanobody-free system, GEM-Halo-FKBP was inserted under the EF1α promoter into Addgene plasmid #129715 after initial insertion of an additional multiple cloning site. To generate a donor plasmid for parallel knock-in of MitoTrap-mCherry-FRB into the AAVS1, MitoTrap-mCherry-FRB was inserted under the EF1α promoter into Addgene plasmid #129719 after initial insertion of an additional multiple cloning site. Two Cas9/sgRNA-expressing plasmids to be used with the AAVS1 donor plasmids were based on #129725^27^ (AAVS1-1 target sgRNA1: ACCCCACAGTGGGGCCACTA GGG and AAVS1-2 target sgRNA2: GTCACCAATCCTGTCCCTAG TGG). The donor plasmids and full sequences will be deposited on Addgene.

### Generation of GEM2-expressing HeLa cell lines via CRISPR/Cas9 genome editing

HeLa cells stably expressing Mito-EGFP or endogenously tagged mEGFP-Ki-67 were electroporated with the two Cas9/sgRNA-expressing plasmids, 5 μg each, and 7.5 μg GEM2 donor plasmid using the Neon Transfection System (Thermo Fisher Scientific) with 3 × 10 ms pulses at 1300 V. Electroporated cells were selected with 0.5 μg/ml puromycin for 2 weeks. To select for the presence of surface protein Thy1.1, trypsinized cells were incubated with 0.4 μg/ml anti-Thy1.1 antibody conjugated with allophycocyanin (APC) in FACS buffer (2% FBS and 0.5 μM EDTA in PBS) for 30 min on ice. Then, APC-positive cells were isolated onto a 96-well plate by FACS. Expanded cells were validated by microscopy to assess cell morphology, GEM2 expression after doxycycline treatment and GEM2-GFP recruitment efficiency upon rapalog treatment. To generate cell lines for nanobody-free GEM2 labelling, HeLa cells were electroporated with four plasmids, 5 μg each: the same two Cas9/sgRNA-expressing plasmids as above, GEM-Halo-FKBP AAVS1 donor plasmid with a puromycin resistance gene and MitoTrap-mCherry-FRB AAVS1 donor plasmid with a blasticidin resistance gene. To select for parallel knock-ins into different alleles of the AAVS1 locus, cells were treated with 0.5 μg/ml puromycin and 6 μg/ml blasticidin (Sigma) for 2 weeks. For FACS, cells were stained with 50 nM Halo-TMR for 30 min. Cells showing strong red fluorescence, likely due to the presence of both mCherry and TMR staining, were isolated onto a 96-well plate. Expanded cells were validated by microscopy to assess cell morphology, GEM2 and MitoTrap expression, and GEM2-GFP coupling efficiency upon rapalog treatment.

### Generation of GEM2-expressing Sum159 cell lines via CRISPR/Cas9 genome editing

Sum159 seipin-sfGFP cells were transfected with the two Cas9/gRNA-expressing plasmids, 0.7 μg each, and 1.05 μg GEM2 donor plasmid as above, using Lipofectamine LTX with Plus reagent. One day later, cells were treated with 1 μg/ml puromycin for 7 days. Thereafter, a single clone was isolated by dilution cloning and this clone was further FACS-sorted for low level GEM2 fluorescence. For FACS sorting, GEM2 expression was induced by 0.2 μg/ml doxycycline for 24 h and GEMs were stained for 1 h with 200 nM Halo-JFX646^28^.

### Transient expression of GEM2 in Nup96 U2OS cells

U2OS cells endogenously expressing Nup96-mEGFP were transfected with the GEM2 donor plasmid using PEI and cultured for 1 day, followed by the treatment with 2 μg/ml doxycycline for 1 day.

### Fluorescence recovery after photobleaching

GEM2 knock-in cells stably expressing Mito-EGFP were stained with 100 nM Halo-TMR for 30 min after the treatment with 2 μg/ml doxycycline for 1 day. Photobleaching was carried out on a Zeiss LSM780 using a Plan-Apochromat 63x/1,4 Oil DIC M27 oil-immersion objective. Bleaching was performed in a 4 μm circular region after initial 10 frames using a laser intensity 160-fold higher than the laser intensity used for image acquisition. Images were acquired every 100 ms for the course of the experiment. Intensities were normalised according to a background region outside the cell as described by Halavatyi and Terjung^29^. A single exponential recovery curve was fitted to the mean of single-normalised intensities over time using FRAPAnalyser (https://github.com/ssgpers/FRAPAnalyser).

### Kinetics assays of GEM recruitment to GFP-tagged proteins

To assess the kinetics of the GEM recruitment to Mito-EGFP, cells were seeded onto LabTek 8-well plates (Thermo Fisher Scientific) and treated with 2 μg/ml doxycycline for 2 days. GEM-Halo-FRB was labelled with Halo-TMR and then treated with rapalog (TAKARA) for 15 min, 30 min, and 60 min. Cells were washed with PBS, then fixed with 3.7% formaldehyde in PBS for 15 min and washed with PBS three times.

To assess the kinetics of GEM2 recruitment to Ki-67 on mitotic chromosomes, cells were seeded onto poly-L-lysine-coated (Sigma) LabTek 8-well plates and treated with 2 mM thymidine (Sigma) and 2 μg/ml doxycycline for 24 hours. Cells were washed with pre-warmed medium three times and cultured in fresh medium supplemented with 2 μg/ml doxycycline and 10 μM S-Trityl-L-cysteine (STLC; Sigma) for 14-16 hours to enrich mitotic cells^30^. Cells were labelled with Halo-TMR and then treated with rapalog for 15, 30 and 60 min. Cells were washed with PBS, then fixed with 3.7% formaldehyde in PBS for 15 min and washed with PBS three times.

To assess the kinetics of GEM2 recruitment to Nup96, transfected cells were stained with Halo-JF646 (Promega) and treated with 0.5 μM rapalog for 1, 6 and 12 h. Cells were washed with PBS, fixed with 3.7% formaldehyde in PBS for 15 min and washed with PBS three times.

To assess the kinetics of GEM2 recruitment to seipin, cells were seeded onto ibidi 8-well plates and treated with 0.2 μg/ml doxycycline for 24 hours. Cells were then treated with 0.5 μM rapalog for 1, 5 and 12 h, 500 μM oleic acid for 1 h and stained with Janelia Fluor X 646 for 1 h. Cells were washed with PBS and then fixed with 4% formaldehyde in PBS for 15 min and washed with PBS three times and quenched with 50 mM NhCl4. Cells were kept in PBS until imaging.

Images were acquired on a Zeiss LSM980 confocal microscope, equipped with Airyscan detector and Plan-Apochromat 63x/1.4 Oil DIC M27 objective. For MitoTrap, Ki-67 and Nup96, single confocal z-planes were acquired and analysed. For seipin, z-stacks covering the whole cell were acquired and analysed. Images were processed with identical settings in each dataset using Zen Blue software (Zeiss). Cell boundaries were manually determined in Fiji. GEMs and target proteins (MitoTrap, Ki-67, Nup96 or seipin) were segmented using ilastik^31^ and the fraction of overlapping areas was analysed using CellProfiler^32^.

### Mitotic chromosome area assay

To assess whether GEM recruitment to Ki-67 affects chromosome dispersion, cells were seeded onto LabTek 8-well plates and treated with 2 μg/ml doxycycline for 2 days. GEMs and DNA were stained with Halo-TMR and SiR-DNA (Spirochrome), respectively. Then, cells were arrested in mitosis with 200 ng/ml of nocodazole for 2 hours, and subsequently treated with 0.5 μM rapalog for 15 min, 30 min, and 60 min. Z-stacks of whole live cells were acquired with 3 μm steps on Zeiss LSM780 using an EC Plan-Neofluar 40x/1.30 Oil DIC M27 oil-immersion objective. Mitotic cells were manually cropped and the centre slice was selected based on the mean intensity of the DNA signal. Mitotic chromosomes were segmented and their ensemble area was analysed using ilastik^31^.

### Importin β binding domain (IBB) import assay

To assess whether GEM recruitment to Nup96 affects nuclear transport, U2OS cells were seeded onto LabTek 8-well plates and transfected with IBB-mCherry and the GEM2 donor plasmid and cultured for 1 day followed by treatment with 1 μg/ml doxycycline for 1 day. Cells were then treated with 0.5 μM rapalog for 1 h, 6 h, and 12 h and GEMs were stained 100 nM of Halo-JF646 (Promega) for 1 h before fixation. Fixed cells were washed with PBS, fixed with 3.7% formaldehyde in PBS for 15 min and then washed with PBS three times. DNA was stained with 0.2 μg/ml Hoechst 33342 for 15 min at RT. Images were acquired by a Zeiss LSM980 confocal microscope with an Airyscan detector and Plan-Apochromat 63x/1.4 Oil DIC M27 objective. Cell boundaries and nuclei were determined manually and by Otsu thresholding in Fiji^33^, respectively. Cytoplasmic segmentations were defined by subtracting the nucleus segmentations from whole cell segmentations. Relative IBB mean intensities were calculated as IBB mean intensity in the nucleus divided by IBB intensity in the cytoplasm.

### Lipid droplet area quantification

To assess whether GEM recruitment to seipin affects the total area of LDs per cell, cells were seeded onto Ibidi 8-well dishes and treated with 0.2 μg/ml doxycycline for 24 hours and 0.5 μM rapalog for the indicated times to induce seipin-GEM tethering. Cells were also treated for 1 h with 500 μM oleic acid to induce LD biogenesis. Cells were then washed with PBS twice, fixed in 4% PFA in PBS for 20 minutes and washed again with PBS twice. Nuclei were stained with Hoechst for 5 min at RT and LDs with 0.2 ug/ml LD540^34^ in PBS for 20 min at RT. Z-stacks of whole cells were acquired with 0.3 μm steps on a Nikon Ti-E widefield microscope with CFI P-Apo DM 60x/1.4 Lambda oil objective. LDs were segmented with Ilastik, and total LD areas per cell were analysed using CellProfiler and Object Analyser as described^15^.

### Fluorescence correlation spectroscopy (FCS)-calibrated imaging

To quantify GFP-tagged protein abundances in living cells, 7.5×10^3^ Mito-EGFP, mEGFP-Ki-67, Nup96-mEGFP and seipin-sfGFP cells were seeded into individual chambers of an 18-well ibidi glass bottom slide in the respective media alongside non-transfected HeLa wild-type cells for estimation of background photon counts, and transfected HeLa wild-type cells with mEGFP for calibration. Prior to imaging, medium was changed to HEPES-based imaging medium (30 mM HEPES pH 7.4, Minimum Essential Eagle medium (Sigma), 10% (v/v) FBS, 1X Minimum Essential Medium non-essential amino acids (Gibco)) containing 100 nM 5-SiR-Hoechst^35^ (gift from Gražvydas Lukinavičius). For Mito-EGFP and Nup96-mEGFP cells, 200 nM 5-SiR-Hoechst (gift from Gražvydas Lukinavičius) and 1 μM Verapamil (Spirochrome) was used. After 1 hour, the imaging medium was supplemented with 500-kDa dextran (Thermo Fisher Scientific) conjugated in-house with Dy481XL (Dyomics) to label the extracellular space.

FCS-calibrated imaging was carried out as previously described^16^ on a Zeiss LSM880 using a C-Apochromat 40× 1.20 W Korr FCS M27 water-immersion objective. Confocal volume estimation was carried out by ten 1-min FCS measurements of 10 nM Atto488 (ATTO-TEC) in water. Background fluorescence and background photon counts were determined by FCS measurements at the nucleus and cytoplasm of non-transfected cells. An experiment-specific calibration line was generated by repeated nucleus and cytoplasm-targeted FCS measurements of WT cells expressing a range of levels of mEGFP. This allowed for the determination of an experiment-specific internal calibration factor with which measured GFP fluorescence intensities could be converted into protein concentrations.

Z-stacks of whole cells were acquired in the GFP, SiR-Hoechst, Dy481XL and transmission channels. A previously established 3D cell segmentation pipeline^36^ was adapted to segment individual cells in large fields of view (Brunner A., Hossain J., Halavatyi A., Morero N. R., Beckwith K. S., Ellenberg J., *in preparation*) and to extract GFP fluorescence intensities for conversion into absolute protein numbers.

### Cryo-ET sample preparation

For all experiments, Au SiO_2_ R1.2/20 Quantifoil grids, 200 mesh, were micropatterned with 30-μm fibronectin circles in the centre of grid squares, as described by Toro-Nahuelpan et al^37^.

For Mito-EGFP, cells seeded in a 6 cm dish were treated with 2 μg/ml doxycycline for 1 day. Then, 2.0×10^5^ cells were seeded onto grids after trypsinization and incubated for 1 h. After cell attachment, grids were transferred to an ibidi 35 mm dishes and further cultured in the medium containing 2 μg/ml of doxycycline for 1 day. Cells were stained with 50 nM Halo-JF646 for 1 h to label GEMs and then treated with 0.5 μM rapalog for 30 min. To verify GEM2 expression and labelling before freezing, fluorescence imaging was performed on a Zeiss Axio Observer microscope, with Plan-Apochromat 63x/1.4 oil objective. Cells were frozen within 30–60 min after rapalog treatment.

For mEGFP-Ki-67, cells were synchronised before seeding by double thymidine block at the G1/S boundary: cells were treated with 2 mM thymidine for 24 h, released and cultured in fresh medium for 8 h, and treated again with 2 mM thymidine for 16–24 h. For seeding, 2.0×10^5^ cells were trypsinized and seeded onto grids in non-arresting fresh medium. The grids were transferred to ibidi 35 mm dishes after cell attachment. 4 h after release, cells were stained with Halo-TMR for 20 min and then DNA was stained with SiR-DNA until freezing. To arrest cells in mitosis, cells were treated with 200 nM nocodazole together with SiR-DNA treatment. Fluorescence montages of the grids were recorded on a LSM780 confocal microscope with an EC Plan-Neofluar 20x/0.50 objective at 37 °C under 5% CO2 at 8.5 h post release. Cells were frozen within 5–10 min of observing mitotic entry at 9–9.5 h post release.

For Nup96-mEGFP, cells transfected with the GEM2 donor plasmid in a 6 cm dish were treated with 2 μg/ml doxycycline for 1 day. Then, 2.0×10^5^ cells were seeded onto grids and incubated for 1 h. The grids were transferred to ibidi 35 mm dishes and treated with 0.5 μM rapalog for 6 hours. Fluorescence imaging was carried out on a LSM780 confocal microscope with an C-Apochromat 63x/1.20 W Corr M27 at 37 °C under 5% CO. Cells were frozen after 7–9 h after rapalog treatment.

For seipin-sfGFP, 4.0×10^5^ cells were seeded onto grids for 20–30 min. After cell attachment, grids were transferred to a new dish for culture and rapalog treatment for 10 h. Cells were treated with Halo-JFX646 for 1.5–2 h and oleic acid for 45–60 min prior to freezing. During JFX646 labelling, montages of the grids were acquired for GEM2 fluorescence on a Zeiss Axio Observer microscope, with Plan-Apochromat 63x 1.4 oil objective.

In all experiments, 3 μL medium was added to the cell side of grids before blotting to reduce cell flattening. Grids were blotted from the back at 37 °C, 90% humidity, and plunge-frozen into liquid ethane at −185 °C on a Leica EM GP2 system, and clipped into cryo-FIB auto-grids.

### Cryo-FIB lamella preparation

Cryo-FIB lamellae were prepared on a 45°-pretilt shuttle in an Aquilos FIB-SEM microscope (Thermo Fisher Scientific) as described using SerialFIB^4^. Before milling, inorganic platinum was deposited by sputter coating (1 kV, 10 mA, 10 Pa, 15–20 s), and organometallic platinum via the Gas Injection System at a working distance of 10.6 mm and injection times of 8–11 s. Cells were rough-milled to 1-μm thickness at a stage tilt of 20° with decreasing ion beam currents (1, 0.5, 0.3 nA) and then thinned all together to a target thickness of 200–250 nm at 50 and 30 pA. Lamellae were thinned at the back at a stage tilt of 21-22°, and finally sputter-coated with platinum (1 kV, 10 mA, 10 Pa, 5–15 s) to reduce beam-induced motion during TEM imaging. Milling progress was assessed by scanning electron microscopy (10 kV, 50 pA).

### Cryo-airyscan imaging of FIB lamellae

For cryogenic on-lamella CLEM, milled grids were loaded onto a Zeiss LSM 900 Airyscan microscope equipped with a Linkam cryo-stage. Using a 5x air objective in widefield mode, an overview image was acquired to localise lamellae. Next, z-stacks with 0.5 μm spacing covering 4–6 μm were acquired in Airyscan mode with 488 and 640 laser lines and Zeiss Plan-Neofluar 100x 0.75 air objective, using a pixel size of 79 nm. In addition, reflection mode images were acquired for each z-plane. For each image, four averages between frames were acquired to increase signal to noise ratio. Z-stacks were 2D Airyscan processed and maximum intensity projections were generated. Subsequent correlation of Airyscan images and low magnification TEM images (lamella maps) was performed in the Icy eC-CLEM plugin^38^ using the lamellae shape and features (such as LDs) visible in the reflection images as guiding marks.

### Cryo-ET image acquisition

Cryo-TEM montages and tilt series were collected on a Titan Krios G3 (Thermo Fisher Scientific) equipped with a Gatan K2 Summit detector and Quantum energy filter, or a Gatan K3 detector and BioQuantum energy filter, using SerialEM^39^. Grids were loaded such that the lamella pre-tilt axis aligns with the microscope stage tilt axis. Images were acquired at a pixel size of 3.370 or 3.425 Å/px at 1.5–4 μm defocus, with an electron dose of 2.0–2.5 e^−^/Å^2^ per image fractionated over 8–10 frames. A dose-symmetric tilt scheme^40^ was used with 2° increments starting from the lamella pre-tilt (±13°) and an effective tilt range of +56° to −56°. Data were collected with a 70 μm objective aperture or Volta phase plate inserted, and 20 eV slit width.

### Cryo-ET data processing

CTF estimation and motion correction were performed in WARP^41^. Dose-weighted motion-corrected images were exported for tomographic reconstruction in IMOD^42^ or AreTomo^43^. To train a DeePiCt neural network^7^, tomograms were binned to a pixel size of 13.48 or 13.70 Å and filtered with the following parameters in EMAN2^44^: filter.lowpass.gauss:cutoff_abs=0.25, filter.highpass.gauss:cutoff_pixels=5, normalize, threshold.clampminmax.nsigma:nsigma=3. Spherical labels of 137 Å radius were generated based on coordinates of 161 manually picked particles in EMAN2. A DeePiCt network of depth 2, with 32 initial features and batch normalisation, was trained with an increasing number of particles and iterative refinement of coordinates via subtomogram averaging (described below). The initial training set contained 161 particles from 39 tomograms, whereas the final training set contained 1284 particles from 71 tomograms. The number of grids and tomograms contributing to this final dataset was as follows: Mito, 6 grids, 12 lamellae, 18 tomograms; Ki-67, 3 grids, 3 lamellae, 9 tomograms; Nup96, 2 grids, 6 lamellae, 20 tomograms; seipin, 5 grids, 9 lamellae, 24 tomograms. Subtomograms at 6.85 Å/px with a box size of 128 px and 3D CTF models were reconstructed in WARP. Averaging was performed in RELION^45^ with 1284 particles, I1 symmetry, using a 60-Å-low-pass-filtered map of the encapsulin scaffold (PDB 6X8M) as reference. Membranes were segmented by tensor voting with TomoSegMemTV^46^ and manual cleaning in Amira (Thermo Fisher Scientific). ER-LD contact sites were defined manually in Amira. The distance between a GEM and its target was estimated based on its refined coordinate and distance to the closest annotated membrane or contact-site pixel using a custom script in Python, subtracting away the radius of a GEM particle (12.5 nm).

### Statistical analyses

Statistical analyses were performed in Graphpad Prism 9.3.1. All Dunn’s tests performed were two-sided and multiplicity-adjusted for multiple comparisons: all time points were compared once against the zero time point.

## Data and code availability

The tomogram of GEM2-labelled Mito-GFP and accompanying mitochondria and GEM annotations presented in Fig. 2a, and the subtomogram average of GEM2 presented in Fig. 2b will be deposited into the EMDB. Raw data corresponding to the full Mito-GFP cryo-ET dataset analysed in Fig. 2c will deposited into EMPIAR. The DeePiCt CNN model for automated GEM detection and Python script for GEM-target distance analyses will be available via GitHub (https://github.com/hermankhfung). CRISPR knock-in donor plasmids generated in this study will be deposited into Addgene. Cell lines generated in this study are available upon request.

## Acknowledgements

This work was supported by the Deutsche Forschungsgemeinschaft (DFG) - SPP2191 Project (419120233 to H.K.H.F., 402723784 to S.C.-H. and J.M.), the European Molecular Biology Laboratory (EMBL) Nuclear Architecture Seed Grant (to H.K.H.F. and Y.H.), the EMBL Interdisciplinary Postdoctoral Program under Marie Skłodowska-Curie Actions COFUND (664726 to H.K.H.F. and Y.H.), the Human Frontier Science Program (CDA00045/2019 to S.C.-H.), Marie Skłodowska-Curie Actions (101028297 to V.T.S.), the Biomedicum Helsinki Foundation (to V.T.S), the Orion Foundation (to V.T.S), and the Boehringer Ingelheim Fonds PhD fellowship (to A.B.). We thank the laboratories of Tobias Walther and Robert Farese for kindly providing endogenous seipin-sfGFP knock-in and seipin knock-out cell lines, the laboratory of Daniel Gerlich for providing endogenous EGFP-Ki-67 knock-in and Ki-67 knock-out cell lines. We thank the EMBL cryo-EM platform, in particular, Wim Hagen, for support in cryo-EM data acquisition, the EMBL Advanced Light Microscopy Facility for support in fluorescence image acquisition and analysis, the EMBL Flow Cytometry Core Facility for support in cell sorting, Thomas Hoffmann and EMBL IT for computational support. We acknowledge the access and services provided by the Imaging Centre at the European Molecular Biology Laboratory (EMBL IC), generously supported by the Boehringer Ingelheim Foundation, and thank Zhengyi Yang for his support. We thank Federico Marotta for advice on tree visualisations.

## Contributions

H.K.H.F., Y.H., V.T.S., C.W.M., S.C.-H. and J.M. conceived this study. H.K.H.F. and Y.H. designed and performed the GEM library screen. Y.H. and V.T.S. performed genome editing and all fluorescence-based kinetic analyses. H.K.H.F., Y.H. and V.T.S. performed cryo-ET imaging with support from A. Babenko and I.Z. H.K.H.F. and V.T.S. analysed the cryo-ET data. A. Brunner performed FCS-calibrated imaging. J.E., C.M., S.C.-H. and J.M. provided supervision. H.K.H.F., Y.H., V.T.S., S.C.-H. and J.M. wrote the manuscript with input from all co-authors. H.K.H.F., Y.H., V.T.S., S.C.-H. and J.M. acquired funding for this project.

**Extended Data Table 1.** GEM library information. Unless otherwise indicated, all constructs were expressed as GEM-Halo-FRB fusions.

| ID | Source organism | Uniprot/RefSeq ID | References |
| --- | --- | --- | --- |
| GEM1 | *Thermotoga maritima* | Q9WZP2 | 47 |
| GEM2 | *Synechococcus elongatus* | Q55032 | 17 |
| GEM3 | *Brevibacterium aurantiacum* | A0A1D7VZ58 | 48 |
| GEM4 | *Paraburkholderia xenovorans* | I2IC15 | 49 |
| GEM5 | *Enterobacteriaceae* | WP_044069342.1 | 49 |
| GEM7 | *Rhodospirillum rubrum* | Q2RVS0 | 50 |
| GEM8 | *Rhodococcus erythropolis* | A0A0E4ACW8 | 51 |
| GEM9 | *Rhodococcus jostii* | Q0SE23 | 52 |
| GEM15 | Synthetic (PCquad) | - | 53 |
| GEM16 | Synthetic (Ico532-1) <sup>†</sup> | - | 54 |
| GEM17 | *Candidatus Scalindua rubra* | A0A1E3XBX5 | 55 |
| GEM22 | *Mycobacterium hassiacum* | K5BEG2 | 56 |
| GEM23 | Synthetic (I3-01) <sup>‡</sup> | - | 57 |

<sup>†</sup> Expressed as an FRB-Halo-GEM fusion
<sup>‡</sup> Expressed as FRB-Halo-GEM and FRB-GEM-Halo fusions

**Extended Data Fig. 1:**
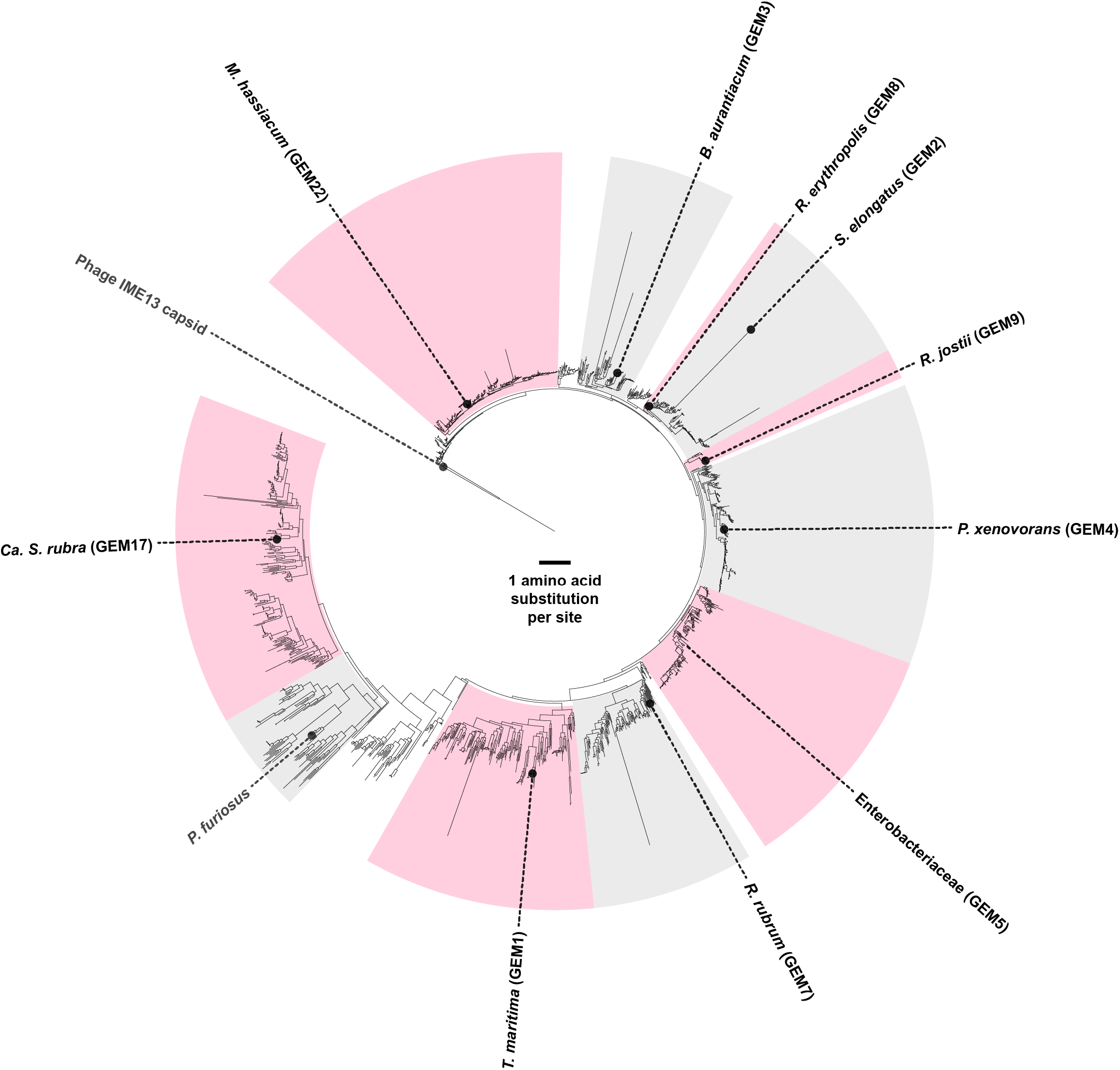
Phylogenetic diversity of selected encapsulins. Maximum likelihood tree of Family 1 encapsulins as identified and constructed by Andreas et al.^10^, with Family 2A encapsulin *Synechococcus elongatus* SrpI (GEM2) and *Stenotrophomonas* phage IME13 capsid protein (outgroup). Scale bar represents amino acid substitution per site. Selected encapsulins, indicated by their GEM IDs in brackets and detailed in Extended Data Table 1, have been shown to form 25-nm-sized particles of triangulation number *T*=1. Also indicated is the *T*=3 40-nm particle-forming encapsulin of *Pyrococcus furiosus*, previously used in budding yeast and HEK293 cells as rheology probes^11^. Clades are shaded in alternating colours up to the most recent common ancestor between annotated sequences.

**Extended Data Fig. 2:**
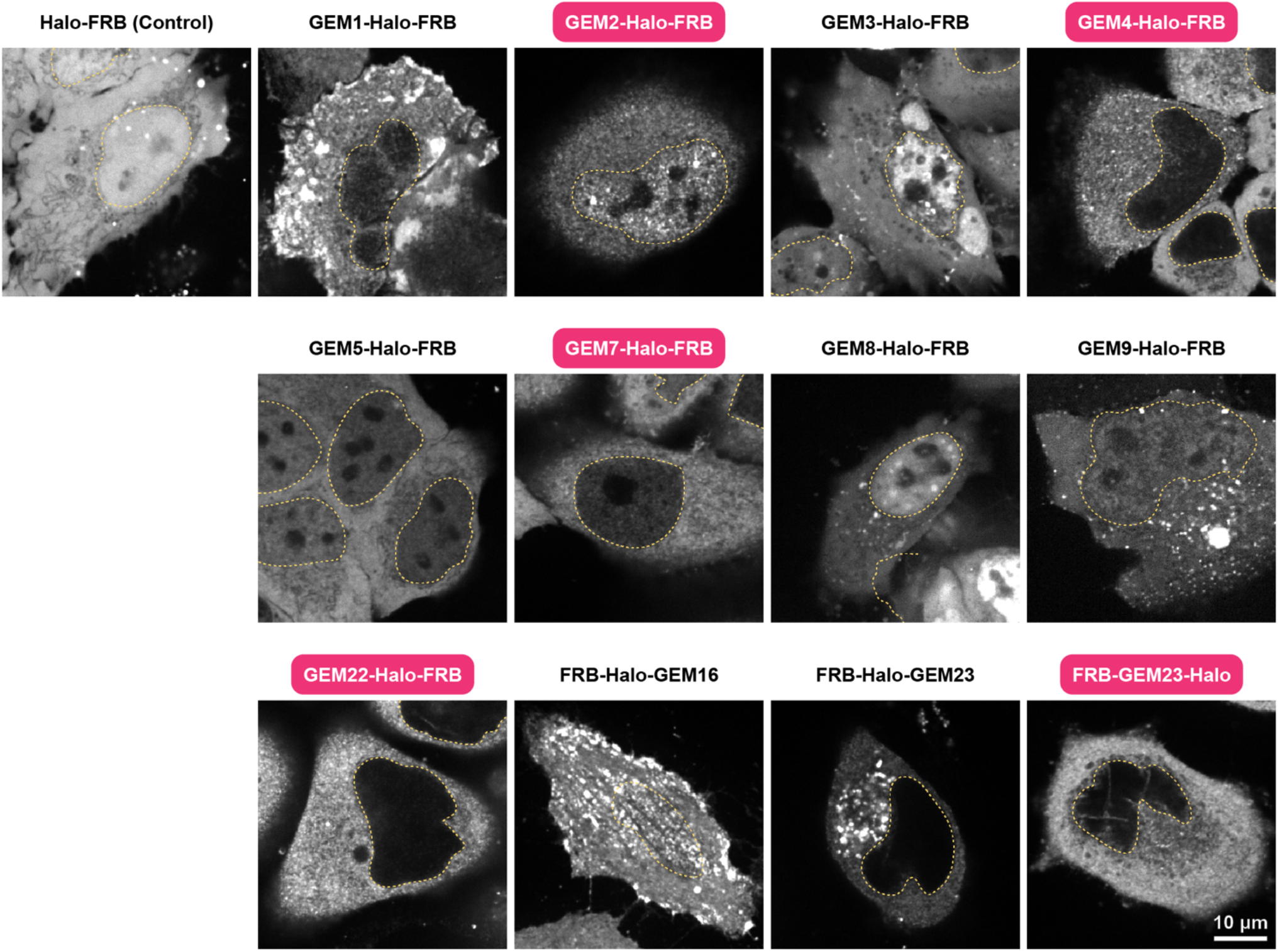
GEM-Halo-FRB fusion expression screen. Constructs were transiently expressed in HeLa cells under a cytomegalovirus (CMV) promoter and labelled with Halo-TMR. Boxed labels in pink indicate constructs that give rise to predominantly uniformly sized fluorescent puncta. All images have the same brightness and contrast setting, except for GEM16 where contrast was enhanced. Orange outlines indicate the cell nucleus based on Hoechst staining (not shown).

**Extended Data Fig. 3:**
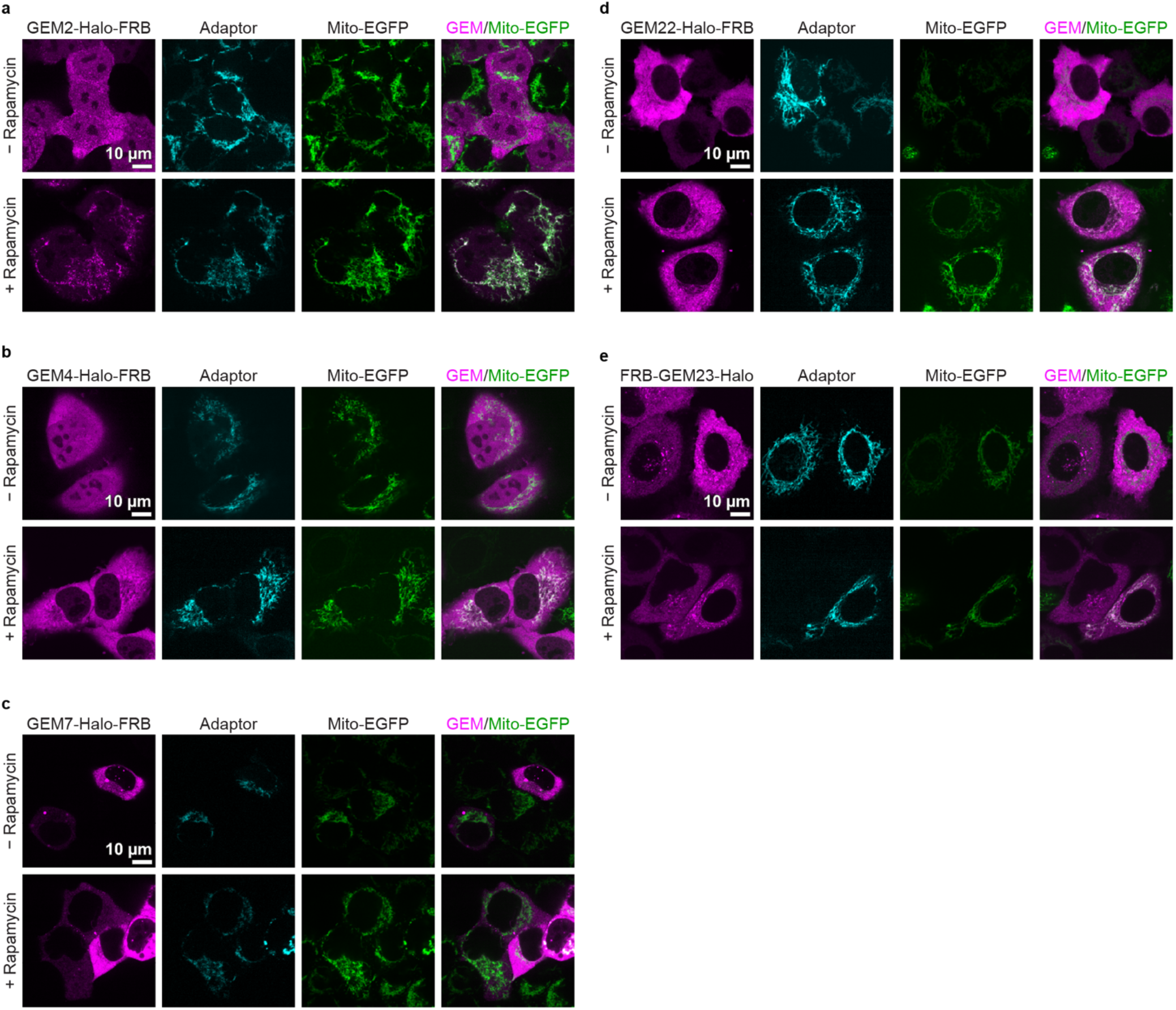
Mito-EGFP GEM coupling assay. **a, b**, HeLa cells stably expressing Mito-EGFP with **a**, GEM2/Adaptor human AAVS1 knock-in or **b**, GEM4/Adaptor human AAVS1 knock-in. Upon 48 h doxycycline induction, GEMs and adaptor proteins were labelled with Halo-TMR and SNAP-SiR, respectively. Cells were treated with rapamycin for 1 h and imaged live. GEM4 showed less efficient labelling of Mito-EGFP than GEM2. **c, d, e**, HeLa cells stably expressing Mito-EGFP and transiently transfected with **c**, GEM7/Adaptor; **d**, GEM22/Adaptor; or **e**, GEM23/Adaptor. Labelling of Mito-EGFP was inefficient in all three cases.

**Extended Data Fig. 4:**
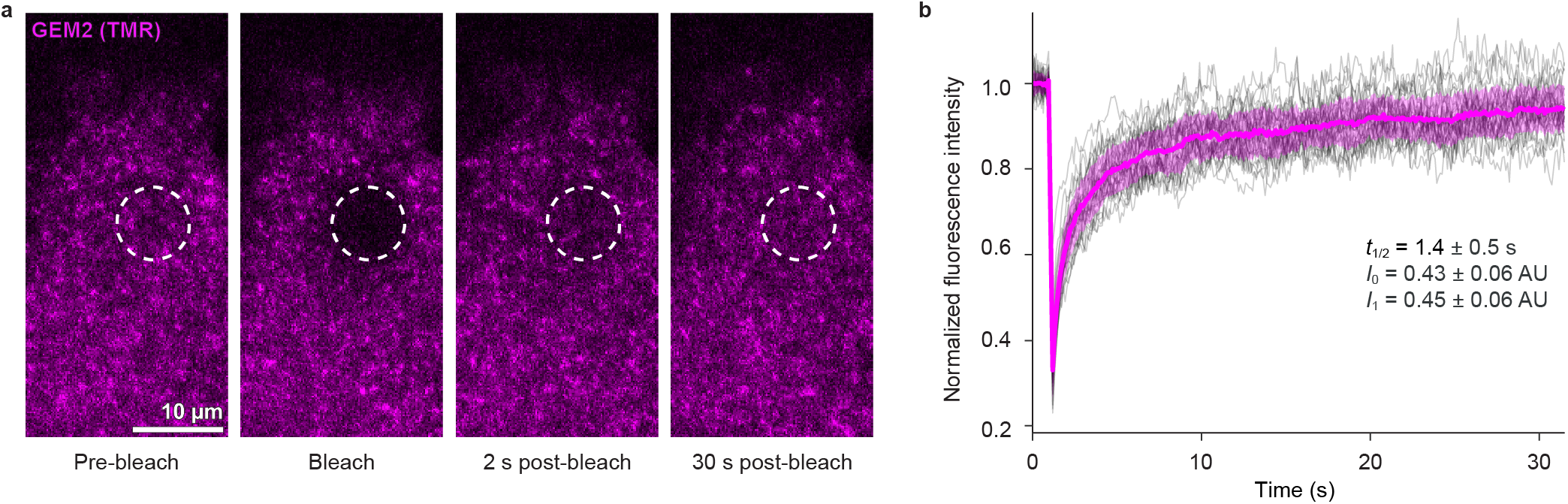
Mobility assay of GEM2. **a**, Fluorescence images of GEMs in HeLa cells stably expressing Mito-EGFP with GEM2/Adaptor knock-in before and after photobleaching, in the absence of rapamycin,. GEMs were labelled with TMR. Dashed circle indicates the photobleached region. **b**, Fluorescence recovery curves. Normalised intensities for individual cells are shown in gray, *n* = 28 cells, 1 experiment. Magenta indicates mean (solid line) ± s.d. (shaded area). A single exponential recovery curve was fitted with *t*_1/2_, *I*_0_, and *I*_1_ representing the recovery half-life, normalised intensity immediately post-bleach, and the dynamic range of recovery.

**Extended Data Fig. 5:**
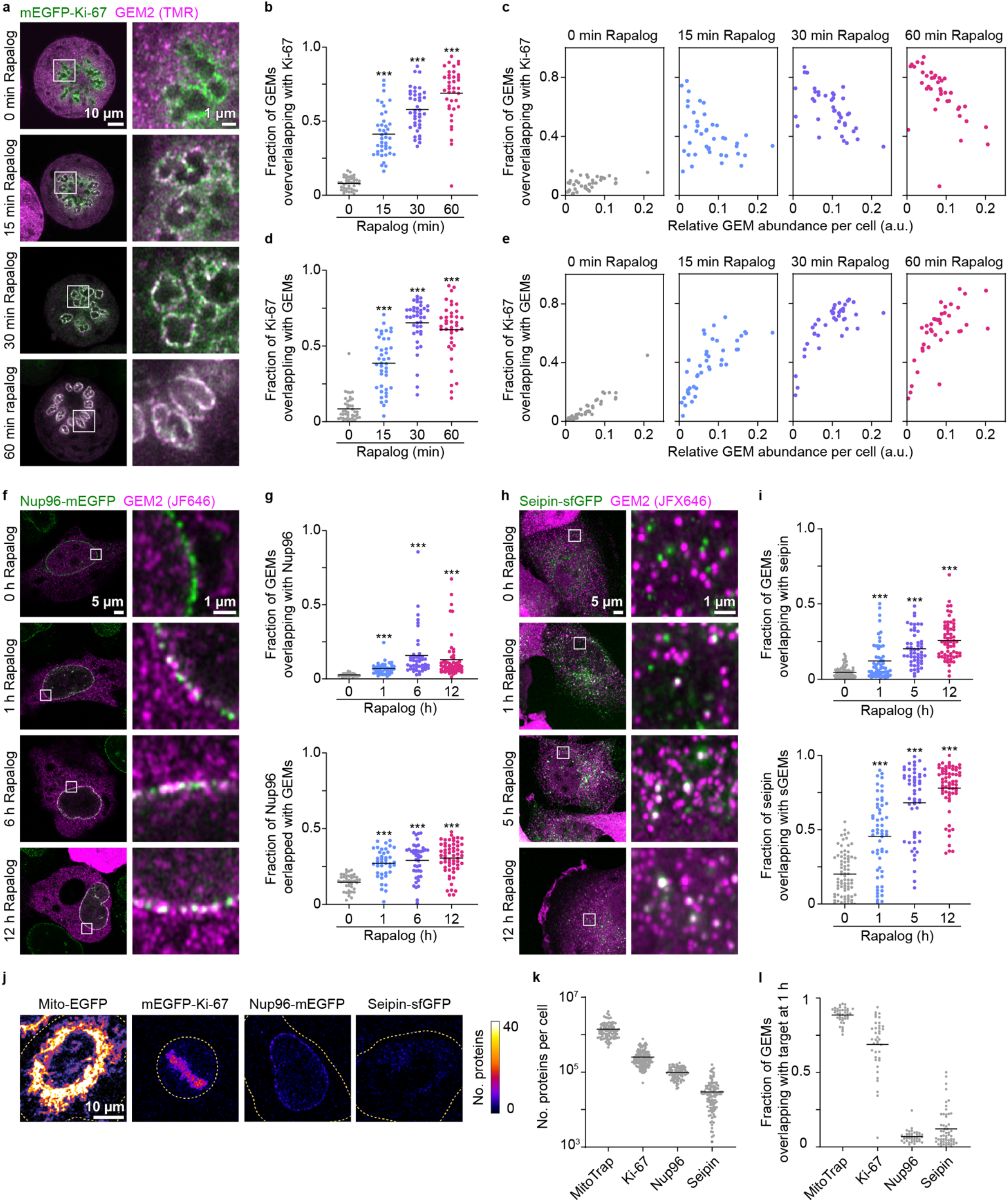
GEM2 recruitment time is dependent on target protein abundance. **a-g**, Kinetic data of GEM2 recruitment to target proteins for Ki-67 (a-e), Nup96 (f, g), and seipin (h, i). GEM2 expression was induced in cells expressing endogenously GFP-tagged proteins and GEM recruitment was induced with rapalog for the indicated durations. **a**, Representative images of GEM recruitment to Ki-67 in mitosis. **b**, Analysis per cell of the fraction of GEM particles overlapping with Ki-67, indicating the extent of GEM particles co-localising with target protein. Lines indicate mean, n = 39-41 cells per group, 2 experiments. P***<0.005, Kruskall-Wallis test followed by Dunn’s test, compared to 0 h rapalog treatment. **c**, Same data plotted as a function of relative GEM abundance per cell, defined as the number of GEM-positive pixels divided by the total cellular area per cell. Longer rapalog treatment times led to more GEMs co-localising with Ki-67. Lower GEM abundance gave rise to a higher fraction of GEMs at the target protein. **d**, Analysis of the fraction of Ki-67 overlapping with GEMs, indicating the completeness of Ki-67 labelling by GEMs. Lines indicate mean, n=39-41 cells per group, 2 experiments. P***<0.005, Kruskall-Wallis test followed by Dunn’s test, compared to 0 h rapalog treatment. **e**, Same data plotted as a function of relative GEM abundance per cell. Longer rapalog treatment times increased labelling of Ki-67 by GEMs. Higher GEM abundance led to more complete coverage of Ki-67. These results demonstrate the importance of tuning GEM expression levels in the labelling experiment. **f**, Representative images of GEM recruitment to Nup96. **g**, Analysis of GEM recruitment to Nup96. Lines indicate mean, n = 41-58 cells per group, 2 experiments. P***<0.005, Kruskall-Wallis test followed by Dunn’s test, compared to 0 h rapalog treatment **h**, Representative images of GEM recruitment to seipin. **i**, Analysis of GEM recruitment to seipin. Lines indicate mean, n = 53-76 cells per group, 2 experiments. P***<0.005, Kruskall-Wallis test followed by Dunn’s test, compared to 0 h rapalog treatment. **j**, Representative images of indicated cell lines. Images were calibrated by FCS to convert GFP fluorescence intensities into absolute protein copy numbers. Yellow dashed lines indicate cell boundaries. **k**, Analysis of h, total cellular protein abundances as determined by FCS-calibrated imaging combined with 3D segmentation. Lines indicate mean, n= 92-148 cells per group, 2 experiments. **l**, Fraction of GEMs overlapping with different target proteins after 1 h rapalog treatment from a-g and Fig. 1c plotted for comparison with k, suggesting a correlation between target protein abundance and GEM2 recruitment efficiency. Lines indicate mean, n = 39-56 cells per group.

**Extended Data Fig. 6:**
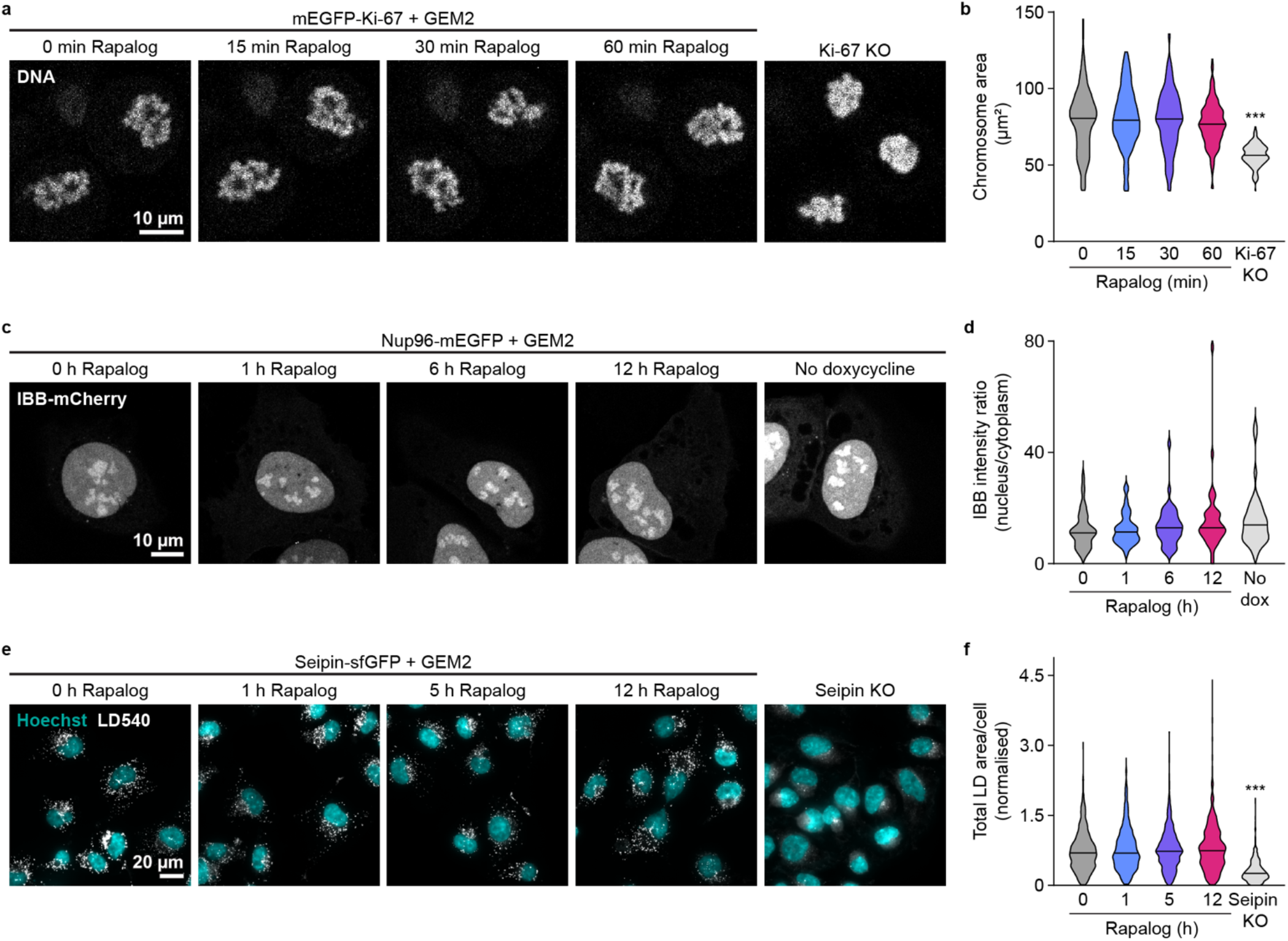
GEM recruitment to the target proteins has little effect on their function. Analysis of target protein function upon induction of GEM2 recruitment for Ki-67 (a, b), Nup96 (c, d), and seipin (e, g). **a**, Representative images of mitotic chromosomes with GEM tethered to Ki-67 after rapalog treatment for the indicated time, in comparison to a Ki-67 knock-out (KO) cell. Ki-67 KO cells serve as a control to indicate that mitotic chromosomes coalesce upon Ki-67 impairment^13^. **b**, Analysis of mitotic chromosome area from a, n=139-188 cells/treatment, 2 experiments. Lines indicate median. P***<0.005, Kruskall-Wallis test followed by Dunn’s test, compared to 0 h rapalog treatment. **c**, Representative images of Importin β binding domain (IBB)-mCherry localization after GEM recruitment to Nup96 at the indicated time treatment with rapalog. Cells not treated with doxycycline do not express GEMs and were used as a control. Impairment of nuclear pore integrity would result in redistribution of IBB to the cytoplasm^58^. **d**, Analysis of relative IBB-mCherry intensities (nucleus/cytoplasm) from c, n=39-42 cells/treatment, 2 experiments. Lines indicate median. P***<0.005, Kruskall-Wallis test followed by Dunn’s test, compared to 0 h rapalog treatment. **e**, Representative images of lipid droplets (LDs) in cells with GEMs recruited to seipin for indicated times and treated with oleic acid for 1 h to induce LD biogenesis. Seipin KO cells serve as a control to indicate total LD area reduction by seipin impairment^15^. **f**, Analysis of LD area per cell from e, n=335-985 cells/treatment, 2 experiments. Lines indicate median. P***<0.005, Kruskall-Wallis test followed by Dunn’s test, compared to 0 h rapalog treatment.

**Extended Data Fig. 7:**
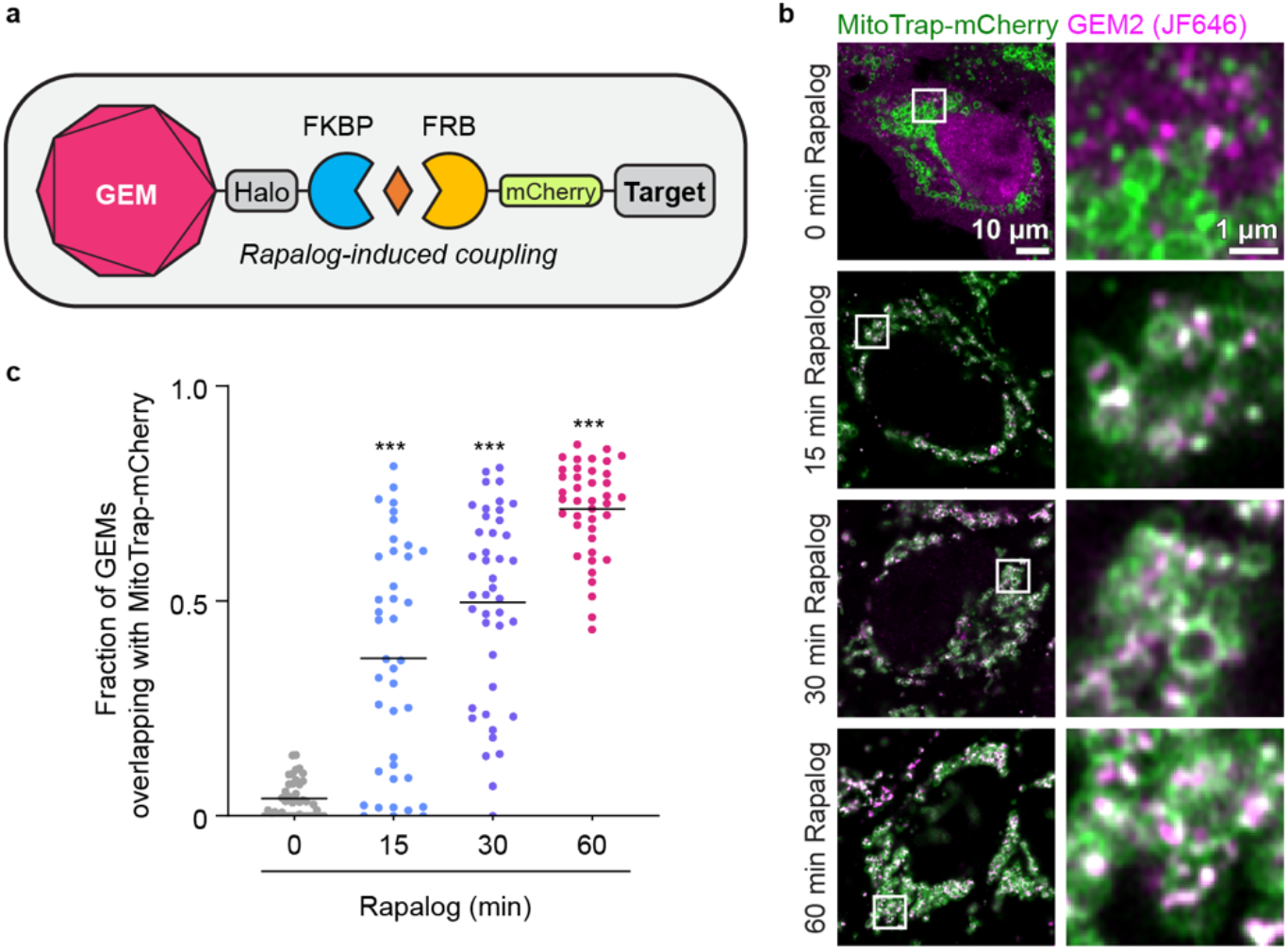
Nanobody-free GEM2 labelling system. **a**, Schematic of the system. **b**, Representative images of GEM2 recruitment to MitoTrap-mCherry upon rapalog treatment by fluorescence microscopy. **c**, Analysis of b. Lines indicate mean, n = 40-42 cells per group, 2 experiments. P***<0.005, Kruskall-Wallis test followed by Dunn’s test, compared to 0 h rapalog treatment.

**Extended Data Table 2.** Cell lines described in this study.

| Cell line name | Comments | Reference | Used in Figs. | Lab ID |
| --- | --- | --- | --- | --- |
| HeLa, wild type | Wild-type cells | Originally from S. Narumiya (Kyoto University, Japan) | Extended Data Fig. 2 | 1 |
| HeLa, Ki-67 KO | Ki-67 knock-out | Published in <sup>13</sup> | Extended Data Fig. 6a,b | 63 |
| HeLa, Mito-EGFP | HeLa stably expressing Mito-EGFP | Generated in this study | Extended Data Fig. 3c–e | 423 |
| HeLa, Mito-EGFP, GEM2-Halo-FRB-IRES-adaptor | Knock-in GEM2-Halo-FRB-IRES-adaptor into AAVS1 locus | Generated in this study | Fig. 1b–d, Fig. 2a–c<br>Extended Data Fig. 3a, 4, 5j–l | 526 |
| HeLa, Mito-EGFP, GEM4-Halo-FRB-IRES-adaptor | Knock-in GEM4-Halo-FRB-IRES-adaptor into AAVS1 locus | Generated in this study | Extended Data Fig. 3b | 526 |
| HeLa, mEGFP-Ki-67, GEM2-Halo-FRB-IRES-adaptor | mEGFP-Ki-67 cells published in <sup>13</sup>, knock-in GEM2-Halo-FRB-IRES-adaptor into AAVS1 locus | Generated in this study | Fig. 1e, Fig. 2b,<br>Extended Data Fig. 5a–e,j–l, 6a,b | 522 |
| HeLa, MitoTrap-mCherry, GEM2-Halo-FKBP | Knock-in GEM2-Halo-FKBP and MitoTrap-mCherry into human AAVS1 locus | Generated in this study | Extended Data Fig. 7 | 539 |
| U2OS, Nup96-mEGFP | Endogenously homozygously labelled Nup96 with C-terminal mEGFP | Published in <sup>22</sup> | Fig. 1f, Fig. 2b,<br>Extended Data Fig. 5f,g,j–l, 6c,d | 466 |
| Sum159, Seipin-sfGFP, GEM2-Halo-FRB-IRES-adaptor | Seipin-sfGFP cells published in <sup>23</sup>, knock-in GEM2-Halo-FRB-IRES-adaptor into AAVS1 locus | Generated in this study | Fig. 1g, Fig. 2b,d,e,<br>Extended Data Fig. 5h–l, 6e,f | - |
| Sum159, Seipin KO | Seipin knock-out | Published in <sup>24</sup> | Extended Data Fig. 6e,f | - |

**Extended Data Table 3.**
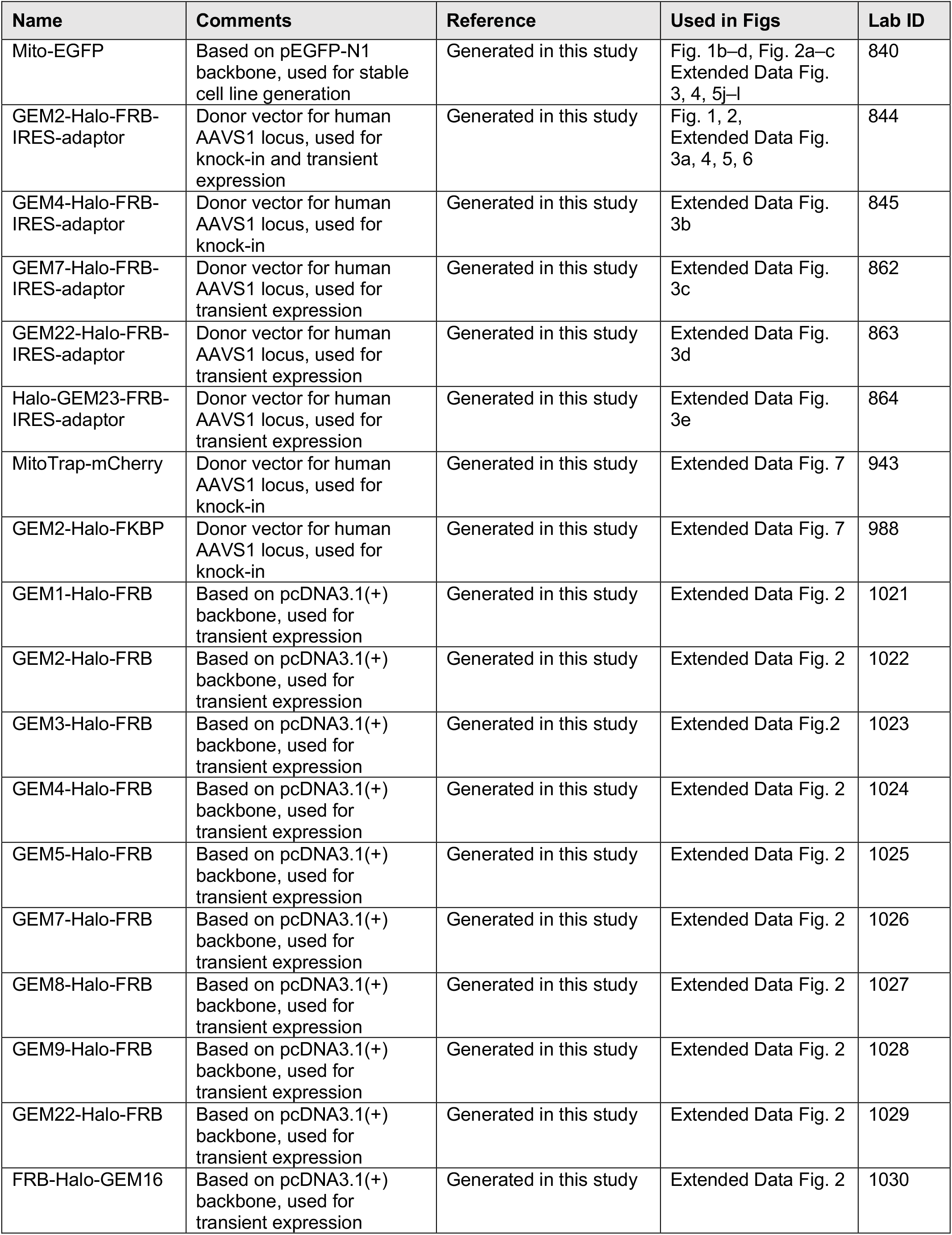
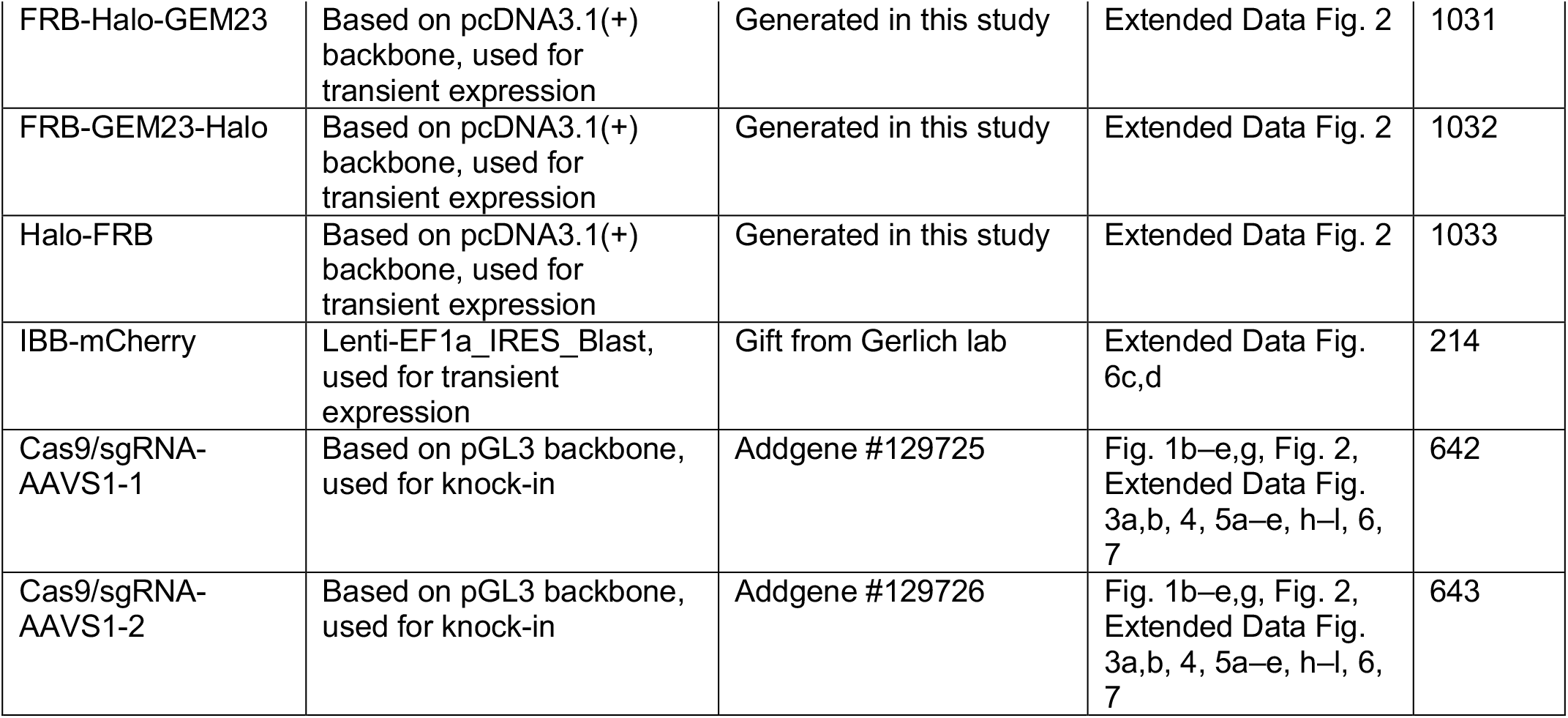
Plasmids described in this study.

| Name | Comments | Reference | Used in Figs | Lab ID |
| --- | --- | --- | --- | --- |
| Mito-EGFP | Based on pEGFP-N1 backbone, used for stable cell line generation | Generated in this study | Fig. 1b–d, Fig. 2a–c<br>Extended Data Fig. 3, 4, 5j–l | 840 |
| GEM2-Halo-FRB-IRES-adaptor | Donor vector for human AAVS1 locus, used for knock-in and transient expression | Generated in this study | Fig. 1, 2,<br>Extended Data Fig. 3a, 4, 5, 6 | 844 |
| GEM4-Halo-FRB-IRES-adaptor | Donor vector for human AAVS1 locus, used for knock-in | Generated in this study | Extended Data Fig. 3b | 845 |
| GEM7-Halo-FRB-IRES-adaptor | Donor vector for human AAVS1 locus, used for transient expression | Generated in this study | Extended Data Fig. 3c | 862 |
| GEM22-Halo-FRB-IRES-adaptor | Donor vector for human AAVS1 locus, used for transient expression | Generated in this study | Extended Data Fig. 3d | 863 |
| Halo-GEM23-FRB-IRES-adaptor | Donor vector for human AAVS1 locus, used for transient expression | Generated in this study | Extended Data Fig. 3e | 864 |
| MitoTrap-mCherry | Donor vector for human AAVS1 locus, used for knock-in | Generated in this study | Extended Data Fig. 7 | 943 |
| GEM2-Halo-FKBP | Donor vector for human AAVS1 locus, used for knock-in | Generated in this study | Extended Data Fig. 7 | 988 |
| GEM1-Halo-FRB | Based on pcDNA3.1(+) backbone, used for transient expression | Generated in this study | Extended Data Fig. 2 | 1021 |
| GEM2-Halo-FRB | Based on pcDNA3.1(+) backbone, used for transient expression | Generated in this study | Extended Data Fig. 2 | 1022 |
| GEM3-Halo-FRB | Based on pcDNA3.1(+) backbone, used for transient expression | Generated in this study | Extended Data Fig. 2 | 1023 |
| GEM4-Halo-FRB | Based on pcDNA3.1(+) backbone, used for transient expression | Generated in this study | Extended Data Fig. 2 | 1024 |
| GEM5-Halo-FRB | Based on pcDNA3.1(+) backbone, used for transient expression | Generated in this study | Extended Data Fig. 2 | 1025 |
| GEM7-Halo-FRB | Based on pcDNA3.1(+) backbone, used for transient expression | Generated in this study | Extended Data Fig. 2 | 1026 |
| GEM8-Halo-FRB | Based on pcDNA3.1(+) backbone, used for transient expression | Generated in this study | Extended Data Fig. 2 | 1027 |
| GEM9-Halo-FRB | Based on pcDNA3.1(+) backbone, used for transient expression | Generated in this study | Extended Data Fig. 2 | 1028 |
| GEM22-Halo-FRB | Based on pcDNA3.1(+) backbone, used for transient expression | Generated in this study | Extended Data Fig. 2 | 1029 |
| FRB-Halo-GEM16 | Based on pcDNA3.1(+) backbone, used for transient expression | Generated in this study | Extended Data Fig. 2 | 1030 |
| FRB-Halo-GEM23 | Based on pcDNA3.1(+) backbone, used for transient expression | Generated in this study | Extended Data Fig. 2 | 1031 |
| FRB-GEM23-Halo | Based on pcDNA3.1(+) backbone, used for transient expression | Generated in this study | Extended Data Fig. 2 | 1032 |
| Halo-FRB | Based on pcDNA3.1(+) backbone, used for transient expression | Generated in this study | Extended Data Fig. 2 | 1033 |
| IBB-mCherry | Lenti-EF1a_IRES_Blast, used for transient expression | Gift from Gerlich lab | Extended Data Fig. 6c,d | 214 |
| Cas9/sgRNA-AAVS1-1 | Based on pGL3 backbone, used for knock-in | Addgene #129725 | Fig. 1b–e,g, Fig. 2, Extended Data Fig. 3a,b, 4, 5a–e, h–l, 6, 7 | 642 |
| Cas9/sgRNA-AAVS1-2 | Based on pGL3 backbone, used for knock-in | Addgene #129726 | Fig. 1b–e,g, Fig. 2, Extended Data Fig. 3a,b, 4, 5a–e, h–l, 6, 7 | 643 |

## Supplementary Information

**Supplementary Video 1. Time-lapse imaging of GEM2 in HeLa cells**. GEM2 was transiently expressed in HeLa cells and labelled with Halo-TMR. Imaging was performed at 115 ms intervals on an Olympus IXplore SpinSR spinning disk confocal microscope using a 100×, 1.35 NA silicone immersion objective.

**Supplementary Video 2. Time-lapse imaging of GEM2 recruitment to Mito-EGFP in HeLa cells**. Cells stably expressing Mito-EGFP with GEM2/Adaptor human AAVS1 knock-in were imaged after 48 h doxycycline induction upon rapamycin treatment. Imaging was performed at 1 min intervals on an Olympus IXplore SpinSR spinning disk confocal microscope using a 100×, 1.35 NA silicone immersion objective. Magenta, GEM2 labelled with Halo-TMR. Green, Mito-EGFP.

**Supplementary Video 3. Tomogram and segmentation of GEM2-labelled Mito-EGFP cells**. Tomogram corresponding to slice shown in Fig 2a. Mitochondrial membrane segmentations, labelled Mito, are displayed in green. GEMs are annotated based on refined coordinates from subtomogram averaging in magenta. MT, microtubule. ER, endoplasmic reticulum. Scale bar, 100 nm.

## References

1. Rigort, A. et al. Focused ion beam micromachining of eukaryotic cells for cryoelectron tomography. Proc. Natl. Acad. Sci. U. S. A. 109, 4449–4454 (2012).

2. Arnold, J. et al. Site-Specific Cryo-focused Ion Beam Sample Preparation Guided by 3D Correlative Microscopy. Biophys. J. 110, 860–869 (2016).

3. Dahlberg, P. D. et al. Cryogenic single-molecule fluorescence annotations for electron tomography reveal in situ organization of key proteins in Caulobacter. Proc. Natl. Acad. Sci. U. S. A. 117, 13937–13944 (2020).

4. Klumpe, S. et al. A modular platform for automated cryo-FIB workflows. Elife 10, e70506 (2021).

5. Böhm, J. et al. Toward detecting and identifying macromolecules in a cellular context: Template matching applied to electron tomograms. Proc. Natl. Acad. Sci. 97, 14245–14250 (2000).

6. Moebel, E. et al. Deep learning improves macromolecule identification in 3D cellular cryo-electron tomograms. Nat. Methods 18, 1386–1394 (2021).

7. Teresa, I. de et al. Convolutional networks for supervised mining of molecular patterns within cellular context. bioRxiv 2022.04.12.488077 (2022) doi:10.1101/2022.04.12.488077.

8. Wang, Q., Mercogliano, C. P. & Löwe, J. A ferritin-based label for cellular electron cryotomography. Structure 19, 147–154 (2011).

9. Silvester, E. et al. DNA origami signposts for identifying proteins on cell membranes by electron cryotomography. Cell 184, 1110–1121.e16 (2021).

10. Andreas, M. P. & Giessen, T. W. Large-scale computational discovery and analysis of virus-derived microbial nanocompartments. Nat. Commun. 12, 1–16 (2021).

11. Delarue, M. et al. mTORC1 controls phase separation and the biophysical properties of the cytoplasm by tuning crowding. Cell 174, 338–349.e20 (2018).

12. Daniel, K. et al. Conditional control of fluorescent protein degradation by an auxin-dependent nanobody. Nat. Commun. 9, 1–13 (2018).

13. Cuylen, S. et al. Ki-67 acts as a biological surfactant to disperse mitotic chromosomes. Nature 535, 308–312 (2016).

14. Mosalaganti, S. et al. AI-based structure prediction empowers integrative structural analysis of human nuclear pores. Science 376, (2022).

15. Salo, V. T. et al. Seipin facilitates triglyceride flow to lipid droplet and counteracts droplet ripening via endoplasmic reticulum contact. Dev. Cell 50, 478–493.e9 (2019).

16. Politi, A. Z. et al. Quantitative mapping of fluorescently tagged cellular proteins using FCS-calibrated four-dimensional imaging. Nat. Protoc. 13, 1445–1464 (2018).

17. Nichols, R. J. et al. Discovery and characterization of a novel family of prokaryotic nanocompartments involved in sulfur metabolism. Elife 10, (2021).

18. Lundin, C. et al. Membrane topology of the human seipin protein. FEBS Lett. 580, 2281–2284 (2006).

19. Sui, X. et al. Cryo–electron microscopy structure of the lipid droplet–formation protein seipin. J. Cell Biol. 217, 4080–4091 (2018).

20. Schmitz, M. H. A. et al. Live-cell imaging RNAi screen identifies PP2A-B55alpha and importin-beta1 as key mitotic exit regulators in human cells. Nat. Cell Biol. 12, 886–893 (2010).

21. Robinson, M. S., Sahlender, D. A. & Foster, S. D. Rapid inactivation of proteins by rapamycin-induced rerouting to mitochondria. Dev. Cell 18, 324–331 (2010).

22. Thevathasan, J. V. et al. Nuclear pores as versatile reference standards for quantitative superresolution microscopy. Nat. Methods 16, 1045–1053 (2019).

23. Chung, J. et al. LDAF1 and Seipin Form a Lipid Droplet Assembly Complex. Dev. Cell 51, 551–563.e7 (2019).

24. Wang, H. et al. Seipin is required for converting nascent to mature lipid droplets. Elife 5, (2016).

25. Kaufman, R. J., Davies, M. V., Wasley, L. C. & Michnick, D. Improved vectors for stable expression of foreign genes in mammalian cells by use of the untranslated leader sequence from EMC virus. Nucleic Acids Res. 19, 4485 (1991).

26. Urlinger, S. et al. Exploring the sequence space for tetracycline-dependent transcriptional activators: Novel mutations yield expanded range and sensitivity. Proc. Natl. Acad. Sci. U. S. A. 97, 7963–7968 (2000).

27. Li, S., Prasanna, X., Salo, V. T., Vattulainen, I. & Ikonen, E. An efficient auxin-inducible degron system with low basal degradation in human cells. Nat. Methods 16, 866–869 (2019).

28. Grimm, J. B. et al. A general method to improve fluorophores using deuterated auxochromes. JACS Au 1, 690–696 (2021).

29. Halavatyi, A. & Terjung, S. FRAP and other photoperturbation techniques. in Standard and Super-Resolution Bioimaging Data Analysis 99–141 (John Wiley & Sons, Ltd, 2017). doi:10.1002/9781119096948.CH5.

30. Skoufias, D. A. et al. S-Trityl-L-cysteine is a reversible, tight binding inhibitor of the human kinesin Eg5 that specifically blocks mitotic progression. J. Biol. Chem. 281, 17559–17569 (2006).

31. Berg, S. et al. ilastik: interactive machine learning for (bio)image analysis. Nat. Methods 16, 1226–1232 (2019).

32. Stirling, D. R. et al. CellProfiler 4: improvements in speed, utility and usability. BMC Bioinformatics 22, 1–11 (2021).

33. Schindelin, J. et al. Fiji: an open-source platform for biological-image analysis. Nat. Methods 9, 676–682 (2012).

34. Spandl, J., White, D. J., Peychl, J. & Thiele, C. Live cell multicolor imaging of lipid droplets with a new dye, LD540. Traffic 10, 1579–1584 (2009).

35. Bucevičius, J., Keller-Findeisen, J., Gilat, T., Hell, S. W. & Lukinavičius, G. Rhodamine–Hoechst positional isomers for highly efficient staining of heterochromatin. Chem. Sci. 10, 1962–1970 (2019).

36. Cai, Y. et al. Experimental and computational framework for a dynamic protein atlas of human cell division. Nature 561, 411–415 (2018).

37. Toro-Nahuelpan, M. et al. Tailoring cryo-electron microscopy grids by photo-micropatterning for in-cell structural studies. Nat. Methods 17, 50–54 (2020).

38. Paul-Gilloteaux, P. et al. eC-CLEM: flexible multidimensional registration software for correlative microscopies. Nat. Methods 14, 102–103 (2017).

39. Mastronarde, D. N. Automated electron microscope tomography using robust prediction of specimen movements. J. Struct. Biol. 152, 36–51 (2005).

40. Hagen, W. J. H., Wan, W. & Briggs, J. A. G. Implementation of a cryo-electron tomography tilt-scheme optimized for high resolution subtomogram averaging. J. Struct. Biol. 197, 191–198 (2017).

41. Tegunov, D. & Cramer, P. Real-time cryo-electron microscopy data preprocessing with Warp. Nat. Methods 16, 1146–1152 (2019).

42. Mastronarde, D. N. & Held, S. R. Automated tilt series alignment and tomographic reconstruction in IMOD. J. Struct. Biol. 197, 102–113 (2017).

43. Zheng, S. et al. AreTomo: An integrated software package for automated marker-free, motion-corrected cryo-electron tomographic alignment and reconstruction. J. Struct. Biol. X 6, 100068 (2022).

44. Tang, G. et al. EMAN2: an extensible image processing suite for electron microscopy. J. Struct. Biol. 157, 38–46 (2007).

45. Kimanius, D., Dong, L., Sharov, G., Nakane, T. & Scheres, S. H. W. New tools for automated cryo-EM single-particle analysis in RELION-4.0. Biochem. J. 478, 4169–4185 (2021).

46. Martinez-Sanchez, A., Garcia, I., Asano, S., Lucic, V. & Fernandez, J. J. Robust membrane detection based on tensor voting for electron tomography. J. Struct. Biol. 186, 49–61 (2014).

47. Sutter, M. et al. Structural basis of enzyme encapsulation into a bacterial nanocompartment. Nat. Struct. Mol. Biol. 15, 939–947 (2008).

48. Putri, R. M. et al. Structural characterization of native and modified encapsulins as nanoplatforms for in vitro catalysis and cellular uptake. ACS Nano 11, 12796–12804 (2017).

49. Giessen, T. W. & Silver, P. A. Widespread distribution of encapsulin nanocompartments reveals functional diversity. Nat. Microbiol. 2, 1–11 (2017).

50. He, D. et al. Structural characterization of encapsulated ferritin provides insight into iron storage in bacterial nanocompartments. Elife 5, (2016).

51. Tamura, A. et al. Packaging guest proteins into the encapsulin nanocompartment from Rhodococcus erythropolis N771. Biotechnol. Bioeng. 112, 13–20 (2015).

52. Rahmanpour, R. & Bugg, T. D. H. Assembly in vitro of Rhodococcus jostii RHA1 encapsulin and peroxidase DypB to form a nanocompartment. FEBS J. 280, 2097–2104 (2013).

53. Lai, Y. T. et al. Designing and defining dynamic protein cage nanoassemblies in solution. Sci. Adv. 2, (2016).

54. Cannon, K. A., Nguyen, V. N., Morgan, C. & Yeates, T. O. Design and characterization of an icosahedral protein cage formed by a double-fusion protein containing three distinct symmetry elements. ACS Synth. Biol. 9, 517–524 (2020).

55. Tracey, J. C. et al. The Discovery of Twenty-Eight New Encapsulin Sequences, Including Three in Anammox Bacteria. Sci. Reports 2019 91 9, 1–11 (2019).

56. Lončar, N., Rozeboom, H. J., Franken, L. E., Stuart, M. C. A. & Fraaije, M. W. Structure of a robust bacterial protein cage and its application as a versatile biocatalytic platform through enzyme encapsulation. Biochem. Biophys. Res. Commun. 529, 548–553 (2020).

57. Hsia, Y. et al. Design of a hyperstable 60-subunit protein icosahedron. Nature 535, 136–139 (2016).

58. Hülsmann, B. B., Labokha, A. A. & Görlich, D. The Permeability of Reconstituted Nuclear Pores Provides Direct Evidence for the Selective Phase Model. Cell 150, 738–751 (2012).

